## Supplementary figures and tables for "Functional analysis in a model sea anemone reveals phylogenetic complexity and a role in cnidocyte discharge of DEG/ENaC ion channels"

**This PDF file includes:**

Legend for Figure S1.

Figures S2 to S6.

Tables S1 to S6.

**Figure S1. Molecular phylogenetic tree of the DEG/ENaC channel superfamily.** The tree was constructed with the IQ-Tree software. See Auxiliary File 1.

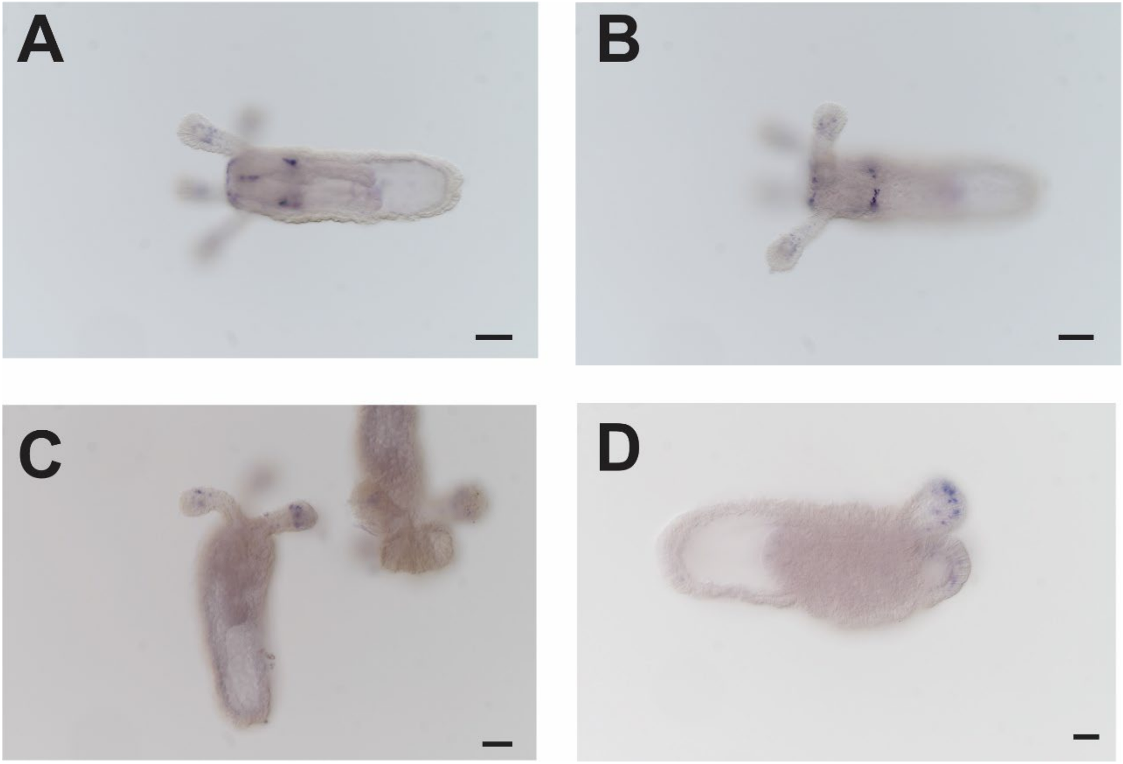


**Figure S2. In situ hybridization of selected NeNaC channel transcripts at the primary polyp stage (9 dpf).** The expression is noticeable by the blue stain (NBT/BCIP crystals). **(A)** NeNaC1; **(B)** NeNaC1; **(C)** NeNaC2; **(D)** NeNaC2. Scale bars: 100 μm.

```
NeNaC2  ------------------------MSLDICDAYIQQETDIGKLYFVNTHILDAAITGCLSLYDLKLIAAVMSCPVAQRHF
NeNaC23 --------------------------------------MSNAPDYRNFIEALHQTLNNPNNRVAVDNRGYALESPLDFYK
NeNaC1  --------MKENDKDEDGGDVKIIIKGSDDKAPQGAGSIYAFCNHCKKDISVVYIKSENNGKELSVAESLTNKWPRSEFP
NeNaC24 -------------------------------------------MALCGCCASKPSPTNSASEDESTGPESKEEILLANQN
NeNaC7  --------------------------------------------------------------------------------
NeNaC6  --------------------------------------------------------------------------------
NeNaC9  --------------------------------------------------------------------------------
NeNaC12 --------------------------------------------------------------------------------
NeNaC10 --------------------------------------------------------------------------------
NeNaC11 --------------------------------------------------------------------------------
NeNaC5  --------------------------------------------------------------------------------
NeNaC3  --------------------------------------------------------------------------------
NeNaC8  --------------------------------------------------------------------------------
NeNaC14 --------------------------------------------------------------------------------
NeNaC15 --------------------------------------------------------------------------------

                                                                     TMD1
NeNaC2  ETEEDKKDEDEDDRAEDPVDENPDDTITVSQMWQDFLHTLTLHGFRFVFERG-PTIRKVLWLAILLFAVGMLMMQSKKSI
NeNaC23 NNWVGDGESVDQKKAQEVDPWNKDEKLTISQHFASLCSSTTMHGISNVFDPSSATLRKGIWSVVFLCSFIFCAYEIGNNV
NeNaC1  ENDYDKCLKDNDYGGCKKLLDGRKKRAKVKELWENFLGGCTLHGFHYCFAGN-PPLRRLIWSLLLLGAFAMFFEKCTESF
NeNaC24 GNASTYENIRKRSAAKRDKKHAEYGKRKMREVFAYYINHCTLHGFHYIFETK-SLFRKIAWFSVLAIAGGFFFKEIKTST
NeNaC7  -------MTSANDDLTNDSRGETNRVGQVRPLWRDFLSRTTLHGTQYACVTK-PLIRRVTWLLLLLGMVGYFGYLFYGNL
NeNaC6  -----MESFNQNTKTDIKDCKEDEKREPQPTMLQDFAGYTTLHGFHFLIHPG-SPFRRFIWLLMLLGCWVALFYQLYNSV
NeNaC9  ------MDPPEERNKEDAREKEHKEENETWNMIREFAGYTTLHGFHFLVDSS-SRWRRRIWFLLLLFCCAMFIYQFVISV
NeNaC12 ----MSYNCKVEDSSDINSNNEYRGGSKTRQLIKEFSGYTTLHGFHFLVDSY-SVTRRVVWTCFIVISLGFLLYQLVNGI
NeNaC10 -MNLFQTKVKPLSRDEAKSDSEKEIEEQRKTLIKNFSSYTTLHGFHFLLDSS-PMPRRVLWTALVVFGLVFFFIQLVMSY
NeNaC11 --MKKDAFETSNQTNFSVFDENEDIRQKRKKLISEFSGYTTLHGLHFLIDSG-SLFRKVFWMILLLVMFTCFFIQLVESY
NeNaC5  ---------MTTVKVANHENPSESVEVPRESLLEAFSSYTTLHGFHFALSST-NRVRQIIWILLLILTSVVLIYQLAYST
NeNaC3  ---------MLKLPCNKEMLRNADPKETALEQINQFLQETTAHGFGRLGATA-GSKWRIYWVMFCLAAYCVFAWQLVGLV
NeNaC8  -------------------METLKPHPSVRAIFKDFSDRTSCHGIGQIGGSQ-SVMWRVSWLMIFLAGLGMVLYQGLTLL
NeNaC14 ---------------------MSRSGPSVRALLRDFSDRTSCHGIGQINGSH-SPTWRIFWLLTFLAGLGMVLFQCITLL
NeNaC15 --------------------MADQPRPSVRALVRDFVDRTTCHGIGQINGSQ-SPLWRVFWVVTFVAGLGMVVHQGVTLF

                                    I
NeNaC2  QKYFDHPITTSVQVEFLE-EIQFPAVTICNFNLFPYYLINGTIGEKVMSI-------------LAPQKYIDNKEEVLFAR
NeNaC23 RYYLTKPVNTVFKIDYVD-EIKFPAVTICNNNPIRKSWAATT-------------------------PY-----------
NeNaC1  INFFDYPFTTTTLLVYDK-RLPFPAISMCNYNDARMSKMNGTLMNEIFVA-------------SKLE-------------
NeNaC24 TQYFKYPFSIMSTVEYPR-TLVFPAVTICDFHDIRQTLVSNAKGEQAIN-------------------------------
NeNaC7  KRYHSHPVEVTVEIETPNDGIGFPAVSVCTNNKYMKSKINMLRN----------------------HSYFHKLGLDIPEC
NeNaC6  DRLLAHRIVMSRGVEQPD-EIDFPAITFCNQNIMRMSKINGTEA----------------------QKYLDELDDVFR--
NeNaC9  NRLRAYKVVFSRGAEQPG-EVDFPAVTICNKNMLRKSLILNTSA----------------------QIYLDEQDADFK--
NeNaC12 KNYNDRGIIMSRSVEEPN-EVDFPAVTICNQNIMKKSLIIGTDA----------------------QRYLDEMNYIKA--
NeNaC10 GKLRARESILAKGVERPM-NVLYPAVTICNQNMMRKSRITGTAA----------------------QRYLDQLDHIKA--
NeNaC11 KRLKEYGSNLSKGVESPE-EVTVPAITICNQNMMRKSLVMGTDA----------------------QKYLDGQDIMKI--
NeNaC5  QRVLEYASMVQVETRNED-SITFPAISICSNNMMQKSKILGKDA----------------------QRYLDLLDQKKE--
NeNaC3  NQYNSKPIKTRTQLKHAQ-KLDFPVVTICNMNVLRASRLPPKLRTKFDEIINNTKKTSSRKSSNSRNAFVDPQDLSFEET
NeNaC8  DTYLNKPTATAVDITYSE-VTNFPSVTICNMNMIKKSQLQHFPQV--KRLVDTFNNMTSSNSSLNSSAFMDTNREKAEKD
NeNaC14 GIYLDKPTATSVDVTYDE-VTNFPAVTICNLNMIKKKNLANFTQS--KKIFDDFEAFVSSNSSMDSSAFLGSKMETVLKD
NeNaC15 GTFLDRPTSTTIDMTYAP-AMDFPAVTICNLNAIRKDHLSQFPDA--DVLLKGFSEAST----PSVTVFLGKDVESFIKQ

                                                                                 II
NeNaC2  SPIPNFLNYAR----KRRRSTGGTIVTDDMLQS----EKD--FGELDEKFDFAEFVRTHGHRIDHM----IKKCRWKS--
NeNaC23 --LPVIMAYNA----NP----GEEAIPIN--------WD----AYNWTGFGFDKLMSSAAHLASEM----IHTCKWKG--
NeNaC1  ---------------------GRNT-------------SH--LQSQLTGELMQRTLKEAAHRLPDM----IKECSWQK--
NeNaC24 ------------------------------------------ISSSE---KLETLARKTYKSFNET----LISCSLRRGV
NeNaC7  AALQNVSGNMTCGQALMCAIVGKYGYIINERCKWALEKIRKIINDSDYAFDIEKFTLKYGHDIKALLT--PRFCTFRG--
NeNaC6  --------------------------------------KD--IKKENVSYDAETFVNKYGHDWENMFENIPYSCMFQR--
NeNaC9  --------------------------------------RS--LQQSNISFDAEEFAKKYGHNITNMLNR-EKGCTFKM--
NeNaC12 --------------------------------D----LGL--VNSTNERLDAEDFVRKYGHTLGEM----MYGCEFKD--
NeNaC10 --------------------------------S----LSR--VNRTNERFETEEMVRLYGHNITDM----LWECNFMN--
NeNaC11 --------------------------------K----LGA--AQVSNESFEVDKMVREKGHLLESM----LFECSFAG--
NeNaC5  --------------------------------D----QWDAISQSFSPDFDIEKAVHQYGLNLSLA----MKSCHYGR--
NeNaC3  K-------------------------------K-----IEILHA--VTTHDNYRELVSAAHQLEDI----LLSCNFNG--
NeNaC8  RL-SLNTKNS-------KNKVILNNVNIDMQRY----IEDKIVQ--YLSMSDTTKLMKAGHVFREL----VFRCVWNG--
NeNaC14 RL-SMDSNDG-------SNSVSLDDTSMDTELY----VEDMLIR--HMAMVDDKDLIEAGHEFDEL----VFRCVWNG--
NeNaC15 HS------------------EAGPNVTLDPELA----FKDAMVE--IFAQSELKKLQMAGHGFEEL----VLGCTWNN--
```

```
                  II                     I
NeNaC2       ---QPCGPE---NFTAV----ITEFG-LCYTFNSGMKGH-----PLLKVQRAGVDYALRLQLSVQQDQYYGSLR-DSSGF
NeNaC23      ---IYCSAA---NFTLD----ATFLG-GCYTFNIDQR---------LMVTGTGMANALHLVLNIQQNEYIGNVR-SGAGF
NeNaC1       --HGKCSWK---NFTSF----KSADGDTCYTFNSGRK-D-----PILSMSNVGEENGLRLVIDTQHSEYYYDVK-N-AGF
NeNaC24      RGARPFNIH---DFKVF----FTAKGQTCYTFNAAMDGK-----KLLEVDNVGPKFGLEIYLNAQHWWFKDDVR-E-SGF
NeNaC7       ---KPCNEE---DFVPV----ITSTS-LCWTFNSGFRGSHGNPAPRKQVTFSGVDFGLTVLLNTRVDENTIGT--SSEGV
NeNaC6       --FFICSAK---NFTSF----LSFTRGLCYTYNSGVNRS-----YVQRVSEAGRNNRLEFHLEAHPEEYYGPYSYEGIGF
NeNaC9       --LYPCSGE---NFTSF----FSFTRGWCYTFNAGA--N-----YIQRVSVAGRETRLKLYLDAKSHEYYGPFSYDGVGF
NeNaC12      ---RRCTAQ---DFIVS----TSFMRGLCYTFNSGRDNS-----SVRRIATPGRLESLILRLNAQPEEYYGAYSYENVGF
NeNaC10      ---KPCSHK---DFAMRY---TSYSRGLCYTFNSGANGS-----PIGQATTSGTRTSLSLRLNAESDEYYGPFSYDATGF
NeNaC11      ---TTCTPE---NFTTS----LSFTRGLCYTFNSGTNNT-----PVFTARAADIRMAFSAMLFSQPEEHYGPFSHRATGF
NeNaC5       --YLKCNPS---HFTTF----KEFRYGLCYTFNSGKRES-----AF-ISHDTGPTSGLSITLDAQPEEYYSLYSSTGTGF
NeNaC3       ---VNCRNSNDPTIPTSWTQTWNDNFGNCYMFNPAQTHN-GEKVDPYSSSIPGESNGLTLQLNIEQNEYLEGIT-EVAGI
NeNaC8       ---FVCNKG---DFLKYWRPFWHWRYGNCYTFNQGVDVN-GTELPSLASSKPGPMYGLTLDLFIDQEQYIIPLS-QEAGV
NeNaC14      ---FTCNKG---GFMKFWRRFWHWRYGNCYIFNQGVDEN-GTLLAHLTSSKPGPMYGLTLDLFIDQEQYLIPLS-QEAGV
NeNaC15      ---IKCNKG---DFLKYWRPFWNFRYGNCYTFNQGMSEK-GVAIKPLTLSNTGPNYGLTLDLFINQEQYIAPYT-QEAGV

                                                          III                         IV  V
NeNaC2       KVMVHDQEEPPLINELGIAIQPGTHTFCGLRKEEMHNLPAPFKTACRDMQ--L----------EGFKKYTKSACLLKCRA
NeNaC23      RLLFREKHEPPSTDRFVIALQPGTQTLIPLTMKKLISLPEPF-GVCQEKNN-L----------KMFDKYSVTACEFECRA
NeNaC1       KVILHDQGETPVK-MQGLSVSPGFTSYMELKRTKVTNLPFPYKTMCGMPE--L----------KYFNSYSKSKCFLDKLT
NeNaC24      RFILHQQDDLPLT-REGFRVSSGYVTYVDMRLEKVKNLPYPFSSDCDGRG--L----------DLYPKYSRNNCYMESLT
NeNaC7       RAVVHEPGEYFSV-DHGVNVMPGAHAAILVHAQKTTTLPLPYKSNCTESK-------------PGLRLYSMEGCVALCAS
NeNaC6       KIAVHDQSYVPNMDQEGYDITAGFYTNVRVKRYKEKSLPHPYKTNCGERK--L----------EYYERYSGSACILECQA
NeNaC9       KIAVHDQNDVPDMDNGGYDISPGYLTTISVKRFKEVSLPPPFPTKCGSRT--L----------EHYERYSTKGCEYECFA
NeNaC12      VLAVHDQAEPPDMELNAYDIPPGFTTNLRIRRFKENSLPEPYPTKCGSRN--L----------SLYKRYSRKACMQECYA
NeNaC10      KLAVHDQNEIPNMDEDAFDTSPGFLTNIRIRREKEINLPSPYRSECGSRD--L----------SNAPKYSMSGCIYECYS
NeNaC11      KIAIHDQSETPDIDLESYDLSPGFATNIRLIREKAKYLPAPYSSNCSSKR--G----------IDGGTYSETGCLTRCYN
NeNaC5       RVIVHDQSEFPWVEKHGWEIPPGFSTNVRLARKEISSLESPYNSNCSRDT--N--------------YASQSYCLVQCYS
NeNaC3       KVSISDQGVLPFPGQQGIRIMPGQSTGIQMTKLQTGRIDPFKNRSCENSN-EMSDKN---LFFGYNMRYSKMACKYSCLN
NeNaC8       KVLLSDQRNIPFPFTHGFTVQPGVSASAGIRQLVIKRIDPFSNGSCYSG-NGLEANSIYHKY--KGMRYSVQGCMSSCLA
NeNaC14      KVLLSDQRNVPFPFTDGFSVQPGVSASVGIRKLVINRIDPFNNGSCYSG-DGLEKENIYSKY--KSLKYSVQGCMSSCLA
NeNaC15      RILLSDQNQIPFPDSDGFTVSPSSSSAVGIKKIFITRIDPFNNGSCYKVTKGLEEGSIYKSVFSHDMGYSVQGCMNSCLA

                  VI VII              VII       VI             V   IV  III
NeNaC2       DYVMKMCKCRSY---DL--KGPAPPC--QPREVKNCVWPAMEIFRNESINC--ECPVPCEITKYQTQLSYAQTPAKHF--
NeNaC23      RLGGKLCGCREMYPSSI--KTEIPVC--LPKAYRDCLNPLLVEISVN-NLC-RGCKNPCNKVTFVPRMSYSQYPANHI--
NeNaC1       QYVVTLCGCRDWFMPG---EGKIPVC--DYETAASCMWKAWAYFEEN-KLD--QCPVACNSVEYSAQLSYARFPAN----
NeNaC24      KYILQQCKCRAWFMAD---VINTSTC--SIKEALDCMWPAWEDFDNA-YNY--TCPVDCEERVYKTRLSSALFLPQKLLP
NeNaC7       QELTRRCGCRPVGLPY---VDAASVC--SFKH-ETCAMDTFGSFDAQ--RC--MCNNACHRTMYNAKVSYARFPDQYIIR
NeNaC6       REFVRKTKCRIIGFPPIKVIRDVPFC--SVLKIETGLTYMYYNWNTN--QC--DCPKPCVEISYSAQMSLLQYPTPSLVR
NeNaC9       KDFVRKFKCKTLGMAPIKEIRNASFC--PVSVVYAALMDFHKNWHLE--LC--DCPKPCETVNYNAQLSTAHYPAPSLLD
NeNaC12      RLIITHCGCRLSGMPPMKDVIEAPFC--TSKEYLDCQLMMPVLLKPS--KC--DCPKRCQHIHYSVQPSLAHYPSKSVIK
NeNaC10      KIIADKCKCRVLGMALN--------------------------VELNPA--MC--DCPKPCRALHYKIQLSLAYFPSDHLWD
NeNaC11      NLMTSQCQCKILGHESDYK-NITGFC--STYQLKACVYEAWMVLRPQ--NC--DCPKPCTSLKYKAQISTSYFPSESLWG
NeNaC5       DMVVKRCGCHMLGMTEE--TGHTPWC--SPQQIKACVYTTSRRFQPN--MC--SCPVRCSRVEFDVQLSSLYYPPDNFWE
NeNaC3       AKTIERCGCTDYNTPELQ-KRNISLCNRLNNAIIDCLNKAYDTFEDG--SCDRECPPSCSEVSFDLTISSAKWPAKSYEK
NeNaC8       NNQFKVCNCTEGKFRA-K-GRP---C--MTEPEVKCLNNISKKYENGSLGCSKSCPQPCTHYSFRRTISQTQWSD-SYEK
NeNaC14      NSEFSICNCTEGKFRV-K-GRP---C--MSESEVKCLNTVNKMYEKGTLGCTRKCPQPCSHFSFRRTISQSQWSE-SYEQ
NeNaC15      NKQREMCNCTEGRFDMMT-GLI---C--QQIEAWRCLNRVNMMYQEGGLKCLEKCPQPCTQNVFSRSMSHAHWAE-EYKK

                                                                             *      TMD2
NeNaC2       ----SEVLARRKHI-----------NKDVMRHYLRDNFLELDVYFEEMQVTLIQQRQAYDQESLFGDIGGQVGLFLGASIL
NeNaC23      ----ADSMAM---------------SMNTTRDFVRDNFLEVEIYFEDIMVEIIEQQEAFSLTSLVGIIGGTLGVFIGASII
NeNaC1       --NYAKMLAKEYGLK---------GSDEENRQYLRDNLIEIKIYYEDLTYFDVQQVPSYDLYSLLGDVGGQIGLFLGASLL
NeNaC24      LTKKYKFLMRPKGIP---------NDTEGAVDFILENYSVINLFFDELRLDTIQQTPAYGFFRLVGDVGGQLGLVLGASVI
NeNaC7       FIQETT------------------SYNYSAEYFRRNLVLVQVGMESLSYEHHRQVPAFPVESLLGAVGGHLGLLLGCSVL
NeNaC6       EIRKSF----------NDT----E---DYINNMRANSVIVSIFYETLLTDVFEEKNDYDISRFGSDLGGNLGLFLGCSLL
NeNaC9       ELSKFPL----EGFVNKSK----E---EYVKFVRDNIVVVEVFYETLLTDVLKEERDYDFNMFASDLGGILGLYLGTSLL
NeNaC12      ELLPSLN----MTEVNSTT----ERINEVNRIIREGHAIIRVFYETLRTEIIKEKPQYTLATLTSDMGGSMGLFLGCSVL
NeNaC10      SIFPVLLNFTELVKVNTTGKSQDEVLLQIQEALRKQIAQVQIYYETLLTDVLEEKPAYGISEFGSDVGGNMGLFLGCSLL
NeNaC11      SLIPFLG-QSSLPFVNLTGKTLEQATAEAQVNVRKSVCMVNVFFETLVTDILEEKPSYDLTMFGADLGGTMGLFLGCSIL
NeNaC5       TISNERS----LTIYAND-------TKKFQEWYRRRVIQLNVFYKELTTEVRKEKEAYTISDLAGDFGGNMGLFLGCSIL
NeNaC3       TVLKTLQT----EYG---------INMTK-EEVFENIAQVHVYYGELDYLLVQETLAYTFMSLLSDIGGQMGMWIGISAL
NeNaC8       TFQRLVRK-SDRGFA---------NKMNDASILRKNFLRVKLYYEELNRETITYSLSYPVENLLGDVGGQLGLWIGVSVI
NeNaC14      TFQKMVMK-SDKGFD---------KKMRDASVLRQNFLRVKLFYEELNTEAITYSRSYTTESFLGDVGGQLGLWIGVSVI
NeNaC15      ALSKFIPN-NGNG-----------SSVDPNELISHNILRVKIYFEELNMETITYKRNYPVESFLGDVGGQLGLWIGVSVI
```

```
            TMD2
NeNaC2   TVLEFLDLLWRILIHKFKKRKNRKVRNV 559
NeNaC23  TVSEFMEFLILIPFRR 494
NeNaC1   TVVEYLDLLGMVAYTSFKYRNG 532
NeNaC24  TIVEIIDLVIMYSIYWIKSKTVPRAATSG---T 506
NeNaC7   TVFEFIDFFIVALASMLRTNVTPNMDRET---KNL 503
NeNaC6   TLVEFFDLGIRWCLGRKDKVQRS 461
NeNaC9   TIAEFLDLGIRWCLRRRSDRSQVRSQAWV---K 470
NeNaC12  TICEFIDLFIQICLERRKRNEVINQK 471
NeNaC10  TFCEFIDLVFMLCLHRHRLRKEAK--------ERQARAAIQDD 470
NeNaC11  TICEFIDLVIILVANGWRKGKARVIDVKE---KPEARP 488
NeNaC5   TIAEFIDLLVVYLVHRHKKRTAKIQ 458
NeNaC3   TCAELVELVCVILANMS---NRSKKIVHISSNIGCFRPAKLGGVEAIEKVGGVKTIEQEVQGGVGIIEQEVQGAVGTIEQ
NeNaC8   TCAEFLKLLVDLAWYLASKMSGKTKTQVQDLNMQ 522
NeNaC14  TCAEFFKLLIDVVWYLARKVHGGPKKTVRDLNMN 520
NeNaC15  TCAEFAKLLIDLVLCVAKKMNRSDKVQSVCIGRGN 510

NeNaC3   EVQGGVGTIEQEMQGRVGTIEQ 578
```

**Figure S3. Sequence alignment of all NeNaCs expressed in *Xenopus laevis* oocytes.** The 15 NeNaCs are shown in the same order as in Fig. 1, putting related NeNaCs together. The conserved N-terminal HG motif, the conserved W residue in TMD1, the selectivity filter in TMD2, and conserved cysteines are shown as white letters on a black background. Disulfide bonds predicted by the crystal structure of cASIC1 are indicated by roman numbers (I-VII). Disulfide bond V is not conserved in closely related NeNaC1 and NeNaC24, and disulfude bond VI is not conserved in closely related NeNaC6 and NeNaC9. NeNaC10 lacks a stretch of about 25 amino acids in the middle of the ECD, including two conserved cysteines. This deletion is due to an alternative exon and likely renders this NeNaC non-functional. Putative positions of TMDs are depicted by black bars. The star indicates the DEG position, close to TMD2.

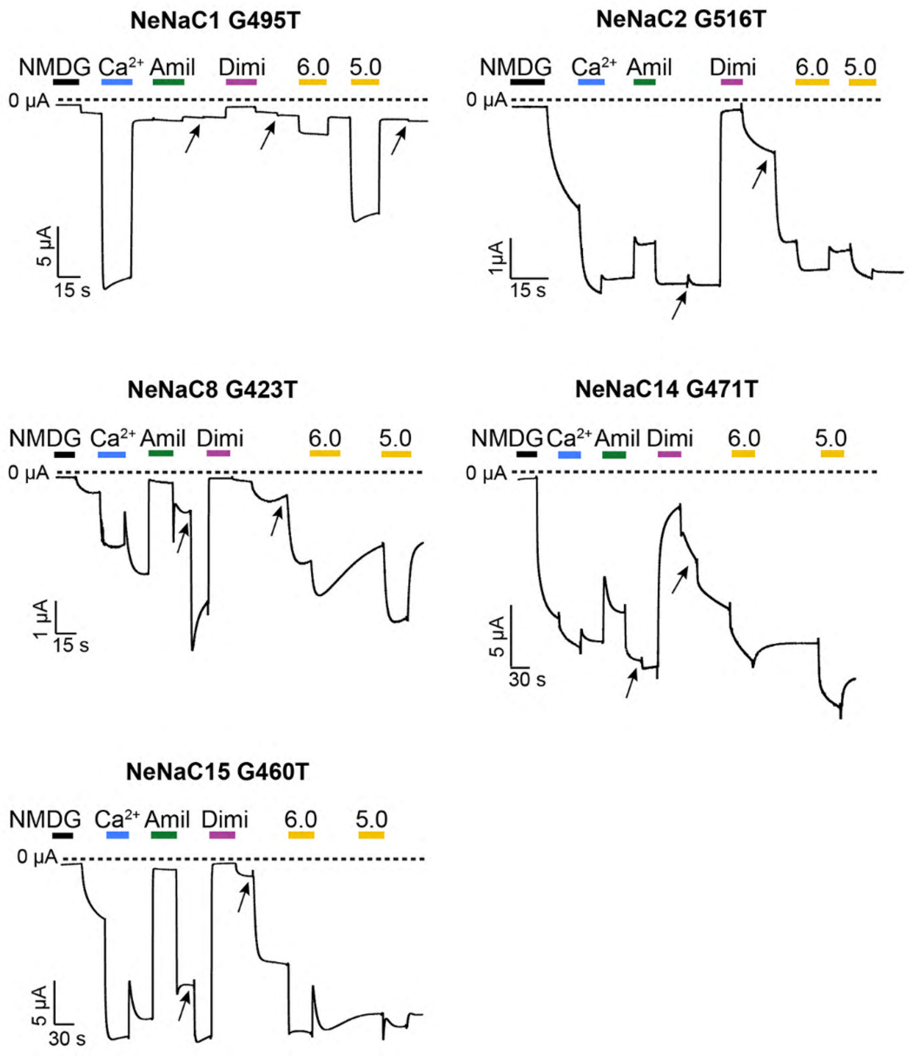


**Figure S4. The DEG mutation constitutively activates some NeNaCs.** Depicted are representative traces of each mutant. They were all activated to a variable extent with the same stimuli: 10 µM $Ca^{2+}$ (blue), pH 6.0 and pH 5.0 (yellow). Moreover, they were blocked by 100 µM amiloride (green) or 10 µM diminazene (dark pink). Our automated solution exchange system exchanges approximately 99% of the bath solution. In some cases, for complete wash-out of blockers, we changed the bath solution twice (indicated by arrows).

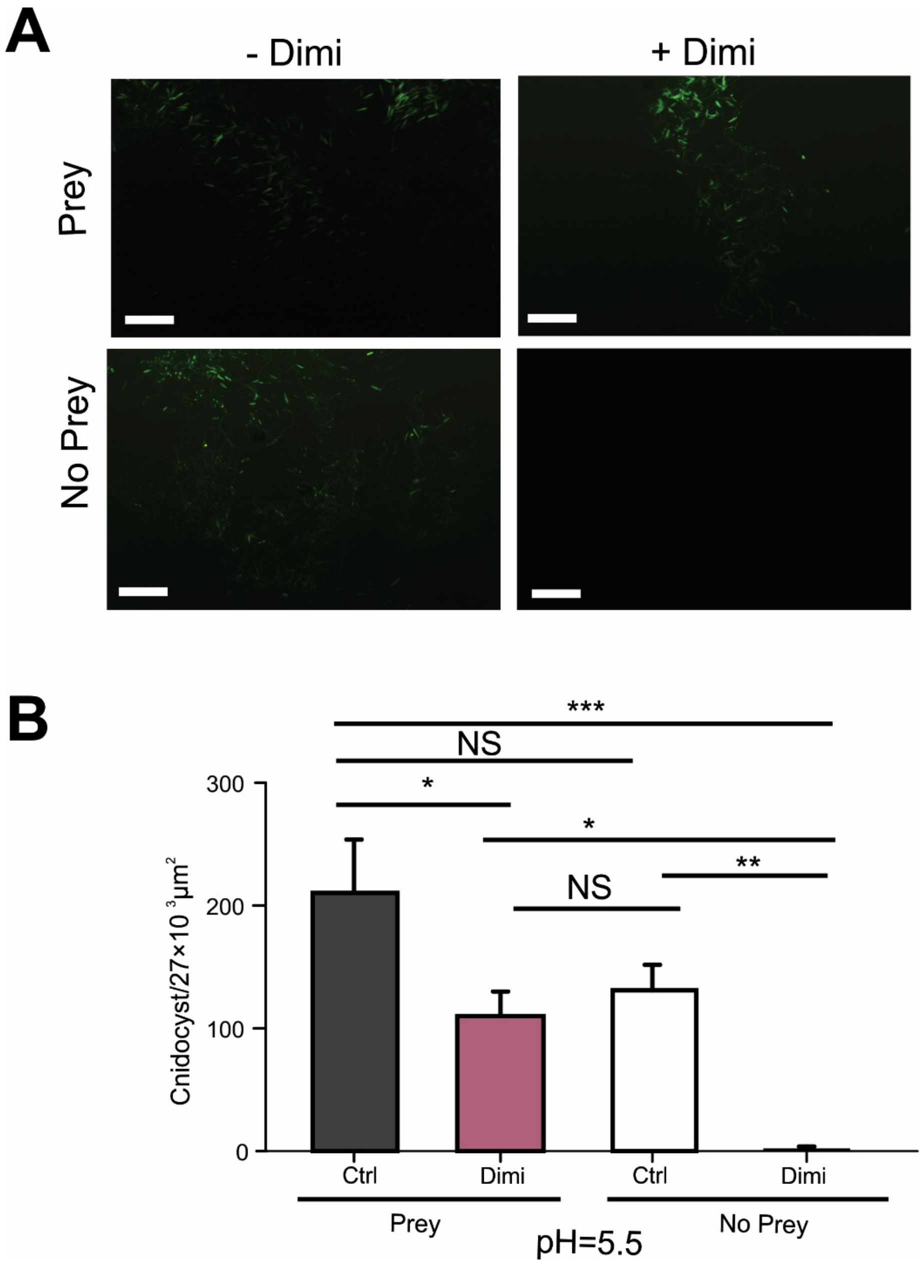


**Figure S5. Cnidocyst discharge at pH 5.5.** (**A**) Pictures of discharged cnidocysts in *NvNCol3::mOrange2* positive organisms, with and without prey extract and with and without diminazene. Scale bar: 100 µm. **(B)** Bar graph showing the number of discharged cnidocysts (mean ± S.E.) at pH 5.5, with and without prey extract and with and without diminazene. One-way ANOVA from 8 individuals per treatment with Tukey post-hoc multiple comparisons test.

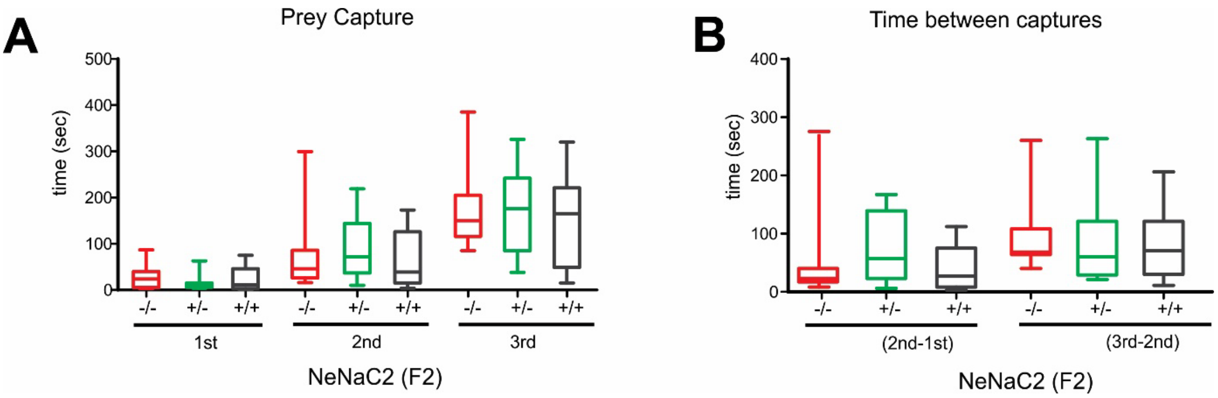


**Figure S6. Prey capture experiment of NeNaC2 F2 progeny from the three genetic pools (NeNaC2$^{(+/+)}$, NeNaC2$^{(+/-)}$ and NeNaC2$^{(-/-)}$).** (A) Box plot with whiskers showing the time in seconds in which NeNaC2 F2 individuals from the three genetic pools capture the first, second and third nauplius at pH 7.2. (B) Box plot with whiskers showing the differences in time between the capture of the second and the first, and the third and the second nauplius at pH 7.2. One-way ANOVA from 15 individuals per treatment with Tukey post-hoc multiple comparisons test for data of **(A)** and (**B)**, and no significant differences were detected.

| NeNaC WT | | | | | | | | | | | | | | | |
|---|---|---|---|---|---|---|---|---|---|---|---|---|---|---|---|
| **Tested Stimuli** | **1** | **2** | **3** | **5** | **6** | **7** | **8** | **9** | **10** | **11** | **12** | **14** | **15** | **23** | **24** |
| pH | - | + | - | - | - | - | - | - | NT | - | - | + | - | - | - |
| amiloride | - | - | NT | NT | NT | - | + | NT | NT | NT | NT | - | NT | - | NT |
| diminazene | - | - | NT | NT | - | - | + | - | NT | NT | NT | - | NT | - | NT |
| low Ca2+ | - | - | - | - | - | - | + | - | - | - | - | - | - | - | - |

| DEG NeNaCs | | | | | | | | | | | | | | |
|---|---|---|---|---|---|---|---|---|---|---|---|---|---|---|
| **Tested Stimuli** | **1 G495T** | **2 G516T** | **3 S464T** | **5 G418T** | **6 S423T** | **7 G404T** | **8 G423T** | **9 S425T** | **10 S419T** | **11 A438T** | **12 S430T** | **14 G471T** | **15 G460T** | **24 G457T** |
| pH | + | + | + | - | - | - | + | - | - | - | - | + | + | - |
| amiloride | + | + | - | - | - | - | + | - | - | - | - | + | + | - |
| diminazene | + | + | - | - | - | - | + | - | - | - | - | + | + | - |

**Table S1. Stimuli tested on NeNaCs and DEG-NeNaCs**. NeNaCs are listed in numerical order from left to right. NeNaCs from clade A are shown on a sky blue background, and NeNaCs from clade B on a reddish purple background. Screening stimuli included acidic pH (mostly 6.0 – 5.0), 100 µM amiloride, 10 µM diminazene and reduction of $[Ca^{2+}]_e$ to 10 µM. A "+" sign indicates that current was elicited or inhibited. A "-" sign indicates that no current was elicited or inhibited. NT = Not Tested.

| Tested Peptides | 1 | 2 | 3 | 5 | 6 | 7 | 8 | 9 | 10 | 11 | 12 | 14 | 15 | 23 | 24 | 1&24 |
|---|---|---|---|---|---|---|---|---|---|---|---|---|---|---|---|---|
| pQGRFa[1] | - | - | - | - | - | NT | - | - | - | - | - | - | NT | NT | - | - |
| VVLRRYa[1] | - | - | - |  | - | NT | - | - | - | - | - | - | NT | NT | - | - |
| WSCSLRPa[1] | - | - | - | - | - | NT | - | - | - | - | - | - | NT | NT | - | - |
| WSCCLRPa[1] | - | - | - | - | - | NT | - | - | - | - | - | - | NT | NT | - | - |
| LVGRWa[1] | - | - | - | - | - | NT | - | - | - | - | - | - | NT | NT | - | - |
| DRTa[1,2] | - | - | - | - | - | NT | - | - | - | - | - | - | NT | NT | - | - |
| pQAGAPGLWa[2] | - | - | - | - | - | NT | - | - | - | - | - | - | NT | NT | - | - |
| pQAGPPGLWa[2] | - | - | - | - | - | NT | - | - | - | - | - | - | NT | NT | - | - |
| 3-L-phenyllactylLRNa[2] | - | - | NT | - | NT | NT | - | NT | NT | NT | NT | - | NT | NT | NT | - |
| 3-L-phenyllactylYRVa[2] | - | - | NT | - | NT | NT | - | NT | NT | NT | NT | - | NT | NT | NT | - |
| pQGLRWa[2] | - | - | NT | - | NT | NT | - | NT | NT | NT | NT | - | NT | NT | NT | - |
| GPRGa[2] | - | - | NT | - | NT | NT | - | NT | NT | NT | NT | - | NT | NT | NT | - |
| pQGRFa[2] | - | - | NT | - | NT | NT | - | NT | NT | NT | NT | - | NT | NT | NT | - |
| pQLFRPa[2] | - | - | NT | - | NT | NT | - | NT | NT | NT | NT | - | NT | NT | NT | - |
| pQLLFRPa[2] | - | - | - | - | - | - | - | - | - | - | - | - | NT | NT | - | - |
| NPPIDLGPAYFHIRa[3] | - | - | - | - | - | - | - | - | - | - | - | - | NT | NT | - | - |
| pQPPIDLSPAAYFHIRa[3] | - | - | - | - | - | - | - | - | - | - | - | - | NT | NT | - | - |
| GPRGGRATEFGPRGa[3] | - | - | - | - | - | - | - | - | - | - | - | - | NT | NT | - | - |
| IPPQGLRFSQWa[3] | - | - | - | - | - | - | - | - | - | - | - | - | NT | NT | - | - |
| MPEQDANPQTRFDa[3] | - | - | - | - | - | - | - | - | - | - | - | - | NT | NT | - | - |
| pQGRFGREDQGRFa[3] | - | - | - | - | - | - | - | - | - | - | - | - | NT | NT | - | - |
| FPPGFHRPa[3] | - | - | - | - | - | - | - | - | - | - | - | - | NT | NT | - | - |
| GPPMIKIPVRHa[3] | - | - | - | - | - | - | - | - | - | - | - | - | NT | NT | - | - |
| pQGRFa[2] | - | - | - | - | - | - | - | - | - | - | - | - | NT | NT | - | - |
| pQAGAPGLWa[2] | - | - | - | - | - | - | - | - | - | - | - | - | NT | NT | - | - |
| pQLLFRPa[2] | - | - | - | - | - | - | - | - | - | - | - | - | NT | NT | - | - |
| pQLFRPa[2] | - | - | - | - | - | - | - | - | - | - | - | - | NT | NT | - | - |

**Table S2. Peptides tested on NeNaCs.** Peptide sequences are listed on the left. Superscript numbers denote references. pQ = pyroglutamate. NeNaCs are listed numerically. NeNaC1 and 24 were also co-injected (right column). NeNaCs from clade A are shown on a sky blue background, and NeNaCs from clade B on a reddish purple background. A "-" sign indicates that no current was elicited or inhibited. NT = Not Tested.

References: [1] Anctil, M. (2009). Chemical transmission in the sea anemone Nematostella vectensis: a genomic perspective. *Comparative Biochemistry and Physiology Part D: Genomics and Proteomics*, *4*(4), 268-289. [2] Koch, T. L., & Grimmelikhuijzen, C. J. (2020). A comparative genomics study of neuropeptide genes in the cnidarian subclasses Hexacorallia and Ceriantharia. *BMC genomics*, *21*(1), 1-21. [3] Hayakawa, E., Watanabe, H., Menschaert, G., Holstein, T. W., Baggerman, G., & Schoofs, L. (2019). A combined strategy of neuropeptide prediction and tandem mass spectrometry identifies evolutionarily conserved ancient neuropeptides in the sea anemone Nematostella vectensis. *PloS one*, *14*(9), e0215185.

| | |
|---|---|
| DNA sequence of NeNaC2 WT | ATGTCGCTTGATATCTGCGATGCCTACATCCAACAGGAAACTGATATCGGAAAACTGTACTTTGTA<br>AATACACACATCTTGGACGCTGCAATAACTGGGTGCTTATCCCTGTATGATTTGAAGTTGATTGCTG<br>CAGTCATGTCGTGTCCAGTTGCACAGAGACATTTCGAAACGGAGGAGGATAAAAAAGATGAAGAC<br>GAGGATGACCGGGCCGAGGATCCTGTTGATGAGAATCCTGATGACACGATCACGGTGTCGCAGAT<br>GTGGCAAGACTTCCTGCACACCCTGACACTACACGGCTTTCGGTTTGTCTTCGAAAGAGGCCCTAC<br>AATTCGCAAGGTGTTATGGCTTGCTATTCTGCTGTTCGCCGTAGGGATGCTGATGATGCAGAGCAA<br>GAAAAGCATACAGAAGTACTTCGACCACCCGATAACGACGAGCGTGCAGGTCGAGTTTCTGGAAG<br>AGATCCAGTTTCCCGCAGTCACAATATGTAACTTTAACCTGTTTCCGTATTATCTTATAAACGGAAC<br>GATCGGCGAAAAGGTGATGTCAATTTTGGCGCCACAGAAATACATCGATAACAAGGAAGAGGTGC<br>TCTTCGCGCGCTCGCCGATACCGAACTTCCTGAATTACGCACGCAAACGGCGTAGGTCAACAGGCG<br>GAACGATAGTAACGGACGACATGCTGCAGAGCGAGAAAGACTTCGGCGAACTCGATGAAAAGTTC<br>GATTTTGCCGAGTTCGTGAGAACGCATGGTCATCGCATCGACCACATGATAAAAAAATGTCGCTGG<br>AAGTCGCAGCCCTGTGGTCCTGAAAACTTTACGGCGGTCATAACAGAATTCGGTCTTTGCTATACCT<br>TCAACTCAGGCATGAAAGGCCACCCTTTACTCAAGGTGCAGAGAGCAGGTGTAGACTACGCCCTTC<br>GGCTGCAGCTCAGCGTTCAGCAGGATCAGTATTATGGCTCCCTGCGCGATTCCTCAGGCTTCAAGG<br>TCATGGTACACGACCAGGAAGAGCCACCACTTATCAACGAGCTCGGCATTGCCATACAACCTGGCA<br>CGCACACGTTCTGCGGCTTGAGAAAAGAAGAGATGCATAATCTCCCAGCGCCGTTCAAAACCGCCT<br>GTCGAGACATGCAGCTAGAAGGCTTCAAGAAATACACCAAGTCAGCATGTCTTTTGAAATGTCGCG<br>CAGACTATGTGATGAAAATGTGCAAGTGTCGCTCTTATGACCTTAAAGGCCCCGCCCCGCCCTGTC<br>AGCCTAGGGAAGTTAAGAACTGCGTTTGGCCCGCAATGGAGATATTCCGCAATGAAAGTATCAACT<br>GCGAGTGTCCAGTCCCTTGTGAGATCACAAAGTACCAAACGCAATTATCTTATGCCCAGACCCCGG<br>CCAAACACTTCTCCGAGGTGCTGGCAAGAAGGAAACACATCAATAAGGATGTCATGAGGCACTAT<br>CTCAGGGATAATTTCTTAGAGCTCGATGTTTACTTCGAGGAGATGCAAGTGACGCTCATTCAGCAG<br>CGACAAGCATATGACCAGGAAAGCTTGTTTGGCGATATTGGTGGTCAAGTAGGGTTGTTTCTTGGA<br>GCAAGCATTCTTACTGTCCTCGAGTTCCTGGACTTGTTATGGAGAATACTCATTCACAAGTTCAAGA<br>AGAGAAAAAAACAGAAAAGTAAGGAATGTATAG |
| DNA sequence of NeNaC2 knock-out line | ATGTCGCTTGATATCTGCGATGCCTACATCCAACAGGAAACTGATATCGGAAAACTGTACTTTGTA<br>AATACACACATCTTGGACGCTGCAATAACTGGGTGCTTATCCCTGTATGATTTGAAGTTGATTGCTG<br>CAGTCATGTCGTGTCCAGTTGCACAGAGACATTTCGAAACGGAGGAGGATAAAAAAGATGAAGAC<br>GAGGATGACCGGGCCGAGGATCCTGTTGATGAGAATCCTGATGACACGATCACGGTGTCGCAGAT<br>GTGGCAAGACTTCCTGCACACCCTGACACTACACGGCTTTCGGTTTGTCTTCGAAAGAGGCCCTAC<br>AATTCGCAAGATGGCTTGCTATTCTGCTGTTCGCCGTAGGGATGCTGATGATGCAGAGCAAGAAAA<br>GCATACAGAAGTACTTCGACCACCCGATAACGACGAGCGTGCAGGTCGAGTTTCTGGAAGAGATC<br>CAGTTTCCCGCAGTCACAATATGTAACTTTAACCTGTTTCCGTATTATCTTATAAACGGAACGATCG<br>GCGAAAAGGTGATGTCAATTTTGGCGCCACAGAAATACATCGATAACAAGGAAGAGGTGCTCTTC<br>GCGCGCTCGCCGATACCGAACTTCCTGAATTACGCACGCAAACGGCGTAGGTCAACAGGCGGAAC<br>GATAGTAACGGACGACATGCTGCAGAGCGAGAAAGACTTCGGCGAACTCGATGAAAAGTTCGATT<br>TTGCCGAGTTCGTGAGAACGCATGGTCATCGCATCGACCACATGATAAAAAAATGTCGCTGGAAGT<br>CGCAGCCCTGTGGTCCTGAAAACTTTACGGCGGTCATAACAGAATTCGGTCTTTGCTATACCTTCAA<br>CTCAGGCATGAAAGGCCACCCTTTACTCAAGGTGCAGAGAGCAGGTGTAGACTACGCCCTTCGGCT<br>GCAGCTCAGCGTTCAGCAGGATCAGTATTATGGCTCCCTGCGCGATTCCTCAGGCTTCAAGGTCAT<br>GGTACACGACCAGGAAGAGCCACCACTTATCAACGAGCTCGGCATTGCCATACAACCTGGCACGC<br>ACACGTTCTGCGGCTTGAGAAAAGAAGAGATGCATAATCTCCCAGCGCCGTTCAAAACCGCCTGTC<br>GAGACATGCAGCTAGAAGGCTTCAAGAAATACACCAAGTCAGCATGTCTTTTGAAATGTCGCGCAG<br>ACTATGTGATGAAAATGTGCAAGTGTCGCTCTTATGACCTTAAAGGCCCCGCCCCGCCCTGTCAGC<br>CTAGGGAAGTTAAGAACTGCGTTTGGCCCGCAATGGAGATATTCCGCAATGAAAGTATCAACTGCG<br>AGTGTCCAGTCCCTTGTGAGATCACAAAGTACCAAACGCAATTATCTTATGCCCAGACCCCGGCCA<br>AACACTTCTCCGAGGTGCTGGCAAGAAGGAAACACATCAATAAGGATGTCATGAGGCACTATCTC<br>AGGGATAATTTCTTAGAGCTCGATGTTTACTTCGAGGAGATGCAAGTGACGCTCATTCAGCAGCGA<br>CAAGCATATGACCAGGAAAGCTTGTTTGGCGATATTGGTGGTCAAGTAGGGTTGTTTCTTGGAGCA<br>AGCATTCTTACTGTCCTCGAGTTCCTGGACTTGTTATGGAGAATACTCATTCACAAGTTCAAGAAGA<br>GAAAAAACAGAAAAGTAAGGAATGTATAG |
| Amino acid sequence of NeNaC2 WT | MSLDICDAYIQQETDIGKLYFVNTHILDAAITGCLSLYDLKLIAAVMSCPVAQRHFETEEDKKDEDEDD<br>RAEDPVDENPDDTITVSQMWQDFLHTLTLHGFRFVFERGPTIRKVLWLAILLFAVGMLMMQSKKSIQK<br>YFDHPITTSVQVEFLEEIQFPAVTICNFNLFPYYLINGTIGEKVMSILAPQKYIDNKEEVLFARSPIPNFLNY<br>ARKRRRSTGGTIVTDDMLQSEKDFGELDEKFDFAEFVRTHGHRIDHMIKKCRWKSQPCGPENFTAVITE<br>FGLCYTFNSGMKGHPLLKVQRAGVDYALRLQLSVQQDQYYGSLRDSSGFKVMVHDQEEPPLINELGIA<br>IQPGTHTFCGLRKEEMHNLPAPFKTACRDMQLEGFKKYTKSACLLKCRADYVMKMCKCRSYDLKGPA<br>PPCQPREVKNCVWPAMEIFRNESINCECPVPCEITKYQTQLSYAQTPAKHFSEVLARRKHINKDVMRHY<br>LRDNFLELDVYFEEMQVTLIQQRQAYDQESLFGDIGGQVGLFLGASILTVLEFLDLLWRILIHKFKKRKN<br>RKVRNV |
| Amino acid sequence of NeNaC2 knock-out line | MSLDICDAYIQQETDIGKLYFVNTHILDAAITGCLSLYDLKLIAAVMSCPVAQRHFETEEDKKDEDEDD<br>RAEDPVDENPDDTITVSQMWQDFLHTLTLHGFRFVFERGPTIRKVAIMACYSAVRRRDADDAEQEKHT<br>EVLRPPDNDERAGRVSGRDPVSRSHNM* |

**Table S3. DNA and Amino acid sequences of NeNaC2 Wild Type Organisms and Knock-out lines (deletion of five nucleotides).** The highlighted green text indicates the start codon, the highlighted yellow text the sgRNA region and the highlighted light blue text the stop codon (DNA). The highlighted magenta text indicates the N-termini, the highlighted navy-blue text indicates the TransMembrane Domains and the highlighted gray text the C-termini (AA).

| NeNaCS | NVE number | UniProtKB | GenBank Number |
|---|---|---|---|
| 1 | NVE20900 | A7RI82 | XP_031568943.1 |
| 2 | NVE13023 | A7RGQ6 | XP_032222736.1 |
| 3* | NVE21425 | A7S2F2 | XP_032239479.1 |
| | NVE9704 | | |
| 4 | NVE574 | A7SA77 | XP_032236192.1 |
| 5 | NVE578 | A7SA83 | XP_032236199.1 |
| 6 | NVE1283 | A7RL38 | XP_032220870.1 |
| 7 | No | A7SH56 | XP_001629025.1 |
| 8 | No | A7SJB5 | XP_032232203.1 |
| 9 | No | A7SST7 | XP_032228649.1 |
| 10 | NVE25684 | A7S8S3 | XP_032236827.1 |
| 11 | NVE6385 | A7SJJ5 | XP_032232107.1 |
| 12 | NVE23003 | A7S4N0 | XP_032238530.1 |
| 13 | No | A7SBZ4 | XP_032232203.1 |
| 14 | NVE15642 | A7SX80 | XP_032226792.1 |
| 15 | NVE10030 | A7SPS3 | XP_032229849.1 |
| 16 | NVE11038 | A7SRD9 | XP_032229236.1 |
| 17 | No | | XP_032233690.1 |
| 18 | NVE3951 | A7SFR1 | XP_032233718.1 |
| 19 | NVE15634 | A7SX71 | XP_001623796.2 |
| 20 | NVE3950 | A7SFR0 | XP_032233720.1 |
| 21 | NVE3518 | A7FS65 | XP_032233957.1 |
| 22 | NVE16828,16829 | A7RWI8 | XP_001636193.2 |
| 23 | NVE12626 | A7STM2 | XP_001625049.2 |
| 24 | NVE20901 | A7RI83 | XP_032222096.1 |
| 25 | NVE24072 | A7RJS8 | XP_032221585.1 |
| 26 | NVE3545 | A7SF92 | XP_032233942.1 |
| 27 | | A7SZH3 | XP_032225907.1 |
| 28 | | A7SZH6 | XP_032224488.1 |
| 29 | | A7SP23 | XP_032230128.1 |

**Table S4. Accession numbers of the 29 NeNaCs from different genomic databases.** NVE models (available at https://figshare.com/articles/dataset/Nematostella_vectensis_transcriptome_and_gene_models_v2_0/807696), UniProt and GenBank. *NeNaC3 has two NVE models.

| **Sequences** |
| --- |
| >Aqueenslandica_XP_011405346.1 |
| >Aqueenslandica_XP_011405345.1 |
| >Pdamicor00010574 |
| >Pdamicor00001790 |
| >Pdamicor00000909 |
| >Actiniatenebrosa17_XP_031566216.1 |
| >Actiniatenebrosa18_XP_031564163.1 |
| >Actiniatenebrosa19_XP_031561615.1 |
| >Amplexidiscus_fenestrater14 |
| >Stylophorapistillata11251 |
| >Stylophorapistillata3268 |
| >Stylophorapistillata14084(1) |
| >Stylophorapistillata104 |
| >Stylophorapistillata4304 |
| >Stylophorapistillata2660 |
| >Mleidy19430 |
| >Mleidy125633 |
| >Mleidy126004 |
| >Mleidy187866 |
| >Mleidy184955 |
| >Scolantuscallimorphus31308 |
| >Scolantuscallimorphus45179 |
| >Scolantuscallimorphus6332 |

**Table S5. Sequences that were removed for the final alignment to construct the molecular phylogenetic tree due to extreme distances or fragmentation.** The sequence of HyNaC1 was not included in the alignment.

| Gene | Forward | Reverse |
|---|---|---|
| NeNaC1 | 5′ATGGCAAAGAGTTGAGCGTGGCCGAGTC | 5′CCGAAAGACCTTGCATTTTGACTGGCGT |
| NeNaC2 | 5′CACATCTTGGACGCTGCAATAACTGGGT | 5′ACTGATCCTGCTGAACGCTGAGCTGCAG |
| NeNaC3 | 5′GGTGGGCCTTGTCAATCAATACAACAGC | 5′ATATTCCGTTTCTGCATCTCAGGGGTGT |
| NeNaC4 | 5′TCAACATGGATGCTATCGTACAGAAGAG | 5′AAGCTCATCGGGTGACAAACCCGTCAGA |
| NeNaC5 | 5′CGAGAAAGTCTTCTCGAAGCCTTCAGCT | 5′GCTTTACCACCATGTCCGAGTAACACTG |
| NeNaC6 | 5′TCCACTTTTTAATTCACCCTGGCTCGCC | 5′CTTGATAGGTGGGAATCCAATAATCCTG |
| NeNaC7 | 5′TCGCCGAGTCACGTGGCTTCTACTTCTG | 5′AAAGGGCTACGCACCCCTCCATTGAGTA |
| NeNaC8 | 5′ATGATATTCCTGGCAGGTCTGGGAATGG | 5′TATGGTCCTCCTGAAAGAGTAATGTGTG |
| NeNaC9 | 5′GTTGGAGGCGCCGCATCTGGTTCCTTTT | 5′CGCGAGGTTTGGGGCAATCACATAACTC |
| NeNaC10 | 5′ATGAACTTGTTCCAGACGAAAGTTAAAC | 5′CTAGTCATCCTGGATTGCTGCCCGGGCT |
| NeNaC11 | 5′CAAGGGCGTCGAGAGTCCAGAAGAAGTC | 5′TTCACGAATGGTAGCGAAGACTGGCCGA |
| NeNaC12 | 5′TTGATTCATACTCCGTAACTCGTCGAGT | 5′CATCTTTCATTGGTGGCATGCCTAACAA |
| NeNaC14 | 5′GCTGTGACCATCTGCAACTTGAACATGA | 5′ACAACCCTGTACGGAATACTTGAGGCTT |
| NeNaC15 | 5′GGACTGGGAATGGTGGTTCATCAAGGCG | 5′GTTTGTTCGCCAGACAGGAGTTCATACA |
| NeNaC16 | 5′GTGGATATTGTTGCTGCTTGTCGCATTT | 5′GCCGGTGGGCAGTTACAAGAAGACATGC |
| NeNaC21 | 5′GCACGGTTTTGCACGACTCGTTGAATCC | 5′GCGTTCTCTGTTAGACCCGATCATTGCC |
| NeNaC22 | 5′CAGCGGCAGCGCTTGTAACCCAATTGAC | 5′TGACCTCGTAGATATTCTCCTCACACGG |
| NeNaC23 | 5′GACCCCTGGAACAAGGATGAGAAACTTA | 5′AACACCAAATGGTTCGGGTAAACTGATC |
| NeNaC24 | 5′CGATCTGCTGCAAAACGAGACAAGAAACA | 5′TGGAGGATGTATTTAGTCAGGCTCTCCA |
| NeNaC25 | 5′TCCCTAAAGACGGCCTCGACTTTCCCGTG | 5′CGTAGATTTGTTCTTTGCACGGGACTGT |

**Table S6. Primers for the generation of amplicons for the ISH experiment.**
