## Supplementary figures and images for "Functional analysis in a model sea anemone reveals phylogenetic complexity and a role in cnidocyte discharge of DEG/ENaC ion channels"

### Supplementary Figure S1

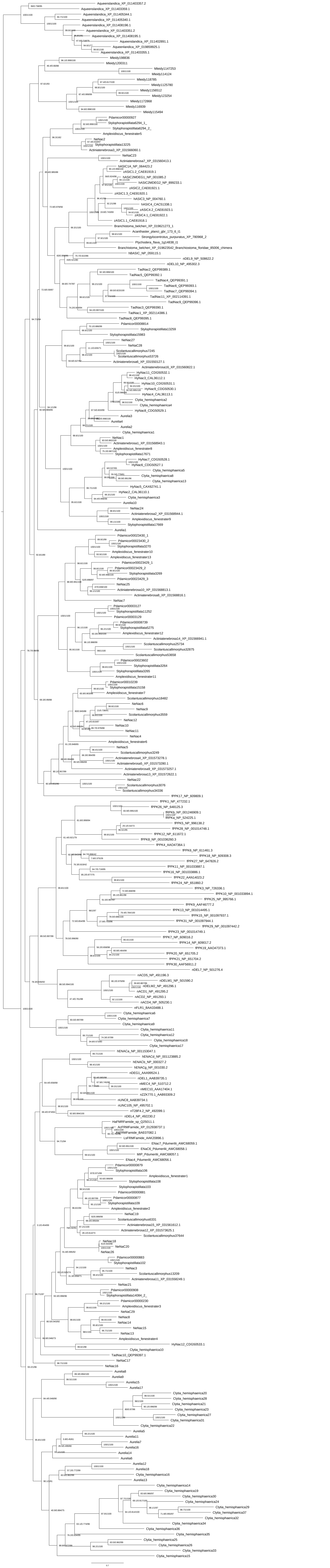
